# Publishing DisGeNET as Nanopublications

**DOI:** 10.1101/010397

**Authors:** Núria Queralt-Rosinach, Tobias Kuhn, Christine Chichester, Michel Dumontier, Ferran Sanz, Laura I. Furlong

## Abstract

The increasing and unprecedented publication rate in the biomedical field is a major bottleneck for discovery in Life Sciences. The scientific community cannot process assertions from biomedical publications and integrate them into the current knowledge at the same rate. The automatic extraction of assertions about entities and their relationships by text-mining the scientific literature is an extended approach to structure up-to-date knowledge. For knowledge integration, the publication of assertions in the Semantic Web is gaining adoption, but it opens new challenges regarding the tracking of the provenance, and how to ensure versioned data linking. Nanopublications are a new way of publishing structured data that consists of an assertion along with its provenance. Trusty URIs is a novel approach to make resources in the Web immutable, and to ensure the unambiguity of the data linking in the (semantic) Web. We present the publication of DisGeNET nanopublications as a new Linked Dataset implemented in combination of the Trusty URIs approach. DisGeNET is a database of human gene-disease associations from expert-curated databases and text-mining the scientific literature. With a series of illustrative queries we demonstrate its utility.

## 1. Introduction

To obtain a deeper understanding of the molecular mechanisms of disease to support drug development and healthcare, biomedical researchers need to explore and query the current knowledge on the complex relationships between genes, proteins, gene variants, pathways, drugs, phenotypes, and environmental factors, among others. Scientific knowledge is initially communicated and gathered as scholarly publications. To ease its access and exploitation, one of the main strategies is to manually extract and curate biomedical statements from the literature and structure them in databases. Due to the increasing size of literature repositories and the number of different, dispersed and isolated databases, there are a lot of efforts devoted to efficiently extracting and providing the most up-to-date data in a way that can be integrated with other statements to facilitate knowledge discovery. Among them, text-mining approaches to unlock relationships between biomedical entities from the literature [1], community-driven publication approaches based on Wiki systems [2-4], or publishing existing databases under the standards of the Semantic Web integrated in a meaningful way to the Linked Open Data cloud (LOD) [5] like UniProt [6], DisGeNET [7], or more ambitious projects like the publication of entire subnetworks such as Bio2RDF [8] and Linked Life Data [9].

DisGeNET is a discovery platform developed in the IBI group designed to enable research on the genetic basis of the pathophysiology of diseases. The platform offers one of the most comprehensive collections of knowledge on human gene-disease associations (GDAs) integrating over 380,000 associations between more than 16,000 genes and 13,000 diseases covering all disease areas. These GDAs are collected from 7 public different databases, which include human and animal model expert-curated databases. Since the literature is a rich, up-to-date source of knowledge on disease genes, we also extracted GDAs derived from MEDLINE by a NLP-based approach named BeFree [10]. All these data are integrated, harmonized and made accessible for exploration and analysis through a Web interface, a Cytoscape plugin [11], and as an RDF Linked Dataset [12], with an open license.

The Semantic Web enables data integration and interoperability, but new challenges emerge when the data are used for the identification and evaluation of scientific hypotheses. These challenges include the tracking of provenance to understand the basis of an assertion and its relation to existing evidence, and the enabling of unambiguous references to immutable pieces of data. To overcome these issues we publish DisGeNET GDAs as a new Linked Dataset using the new nanopublication approach [13] and the Trusty URI technique [14]. The nanopublication approach is a community-driven effort that proposes a publishing model for minimal scientific statements along with their provenance and associated context. A nanopublication is formally structured using Semantic Web technologies (W3C’s Resource Description Framework (RDF) and the Web Ontology Language (OWL)), specifically, using RDF named graphs following the nanopublication schema. Consequently, standard RDF technologies such as triple stores and SPARQL query engines can be used to deal with nanopublications. In addition, we identify nanopublications using Trusty URIs, which is a novel approach to make resources in the Web verifiable, immutable, and permanent, consisting of the integration of cryptographic hash values (representing the content of a nanopublication) in Uniform Resource Identifiers (URIs).

In this paper, we present the DisGeNET Nanopublications Linked Dataset as an alternative way to mine the GDAs contained in DisGeNET. The nanopublication conversion of DisGeNET aims to extend and complement the capabilities of the existing DisGeNET RDF Linked Dataset, to enrich the number of GDAs assertions in the Web of data, to foster the publication, aggregation, mining, discoverability of these assertions, and to support the generation of evidence-based powerful and new hypothesis in the biomedical field.

## 2. Gene-disease Associations in DisGeNET

In order to cover different aspects of the genedisease relation, DisGeNET GDA content is imported from various types of sources ranging from human and animal model expert-curated databases to literature-based datasets obtained by different text-mining approaches. DisGeNET provides an evidence classification according to the level and type of curation in these original databases that enable users to rapidly assess the quality of the specific GDA. The Dis-GeNET evidence classes are: “CURATED” for human GDA evidence reviewed by experts, “PREDICTED” for inferred human GDA from animal model’s GDA evidence reviewed by experts, and “LITERATURE” for automatic extraction of human GDA from the literature (see DisGeNET coverage in Table 1). It is important to point out that DisGeNET not only aggregates GDAs statements from different sources, but integrates them in a uniform way accompanied with context annotation. Remarkably, in DisGeNET, the provenance and evidence is well described and fine-grained for each statement in order to keep them tracked after integration. DisGeNET is represented in various structured data model conventions: as a relational database, as an RDF Linked Dataset, and now as a nanopublication Linked Dataset.

**Tabel 1.**
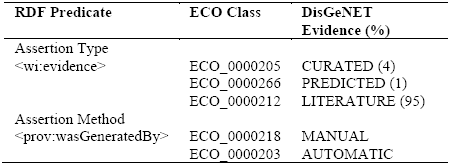
The DisGeNET evidence classification and coverage in the nanopublication dataset.

### 2.1 RDF Dataset Description

The source of data for the DisGeNET nanopublications set is the RDF Linked Dataset version of DisGeNET. There are three main components in the RDF dataset: GDA content, provenance description of the RDF dataset, and linksets to other Linked Datasets. Each of these components is described separately in the following sections.

#### 2.1.1 GDA Content

The current RDF representation is centered on the GDA concept, and different information around GDA, such as the gene and disease involved identified by NCBI Gene ID and UMLS CUI, respectively, and the type of association is represented. Also, the gene and the disease have different annotated attributes (see the schema at http://rdf.disgenet.org/). Entities and properties are semantically annotated using standard ontologies such as the NCI Thesaurus (NCIt) [15], and resources identified by using de-referenceable URIs. GDAs are integrated using the DisGeNET association type Ontology, which is an ontology developed in the IBI group to fill the gap in formal semantics for the definition of types of associations between a gene and a disease in biological databases. This ontology is based on the description of the GDAs association type in the original databases. The DisGeNET ontology is integrated into the Sematicscience Integrated Ontology (SIO) [16], which is an OWL ontology that provides essential types and relations for the rich description of objects, processes and their attributes, so that GDAs in RDF are semantically harmonized using SIO classes. A normalized de-referenceable URI scheme for the identification of GDA entities is implemented using “http://rdf.disgenet.org/” as namespace and a unique identifier built based on unique attributes.

#### 2.1.2 Provenance

A full provenance description of the RDF Linked Dataset is provided using the Vocabulary of Interlinked Datasets (VoID), an RDF Schema W3C recommended vocabulary for expressing metadata about RDF datasets [17]. This description includes the provenance of DisGeNET relational database, primary databases, and the BeFree text-mining approach. The type of curation and level of evidence of each original database are also tracked and annotated. Each data instance in DisGeNET is explicitly referenced to this provenance description in order to granulate and trace back the provenance to the instance level (Access to the VoID.ttl file description at http://rdf.disgenet.org/Website).

#### 2.1.3 Linksets

In addition, linkouts to the LOD are set in order to both enrich DisGeNET GDAs annotations with external Semantic Web resources, and to extend the current GDAs content of the network of bio-entities embedded in the Web of data. Specifically, a total number of 4,962,315 linksets to the LOD through Bio2RDF, LinkedLifeData subnetworks projects among others exists in the current version. All entities linked are related using the same SKOS [18] predicate *skos:exactMatch*. Other linkset statistics between entities can be found at the DisGeNET DataHub site in *thedatahub.org* [19].

## 3. DisGeNET Nanopublication Dataset

A DisGeNET nanopublication is modeled by 4 named graphs: Head, Assertion, Provenance and publicationInfo (Figure 1). The Head graph contains statements regarding the formal nanopublication structure. Basically, the linking between graph URIs in the nanopublication. The assertion graph contains the minimal description for a specific GDA assertion. The provenance graph includes provenance, evidence and attribution statements that were easily implemented thanks to the VoID description of the RDF dataset. Finally, the publication information graph includes all the metadata information regarding the nanopublication itself. We also include in this graph a description of the general topic of the nanopublication to enhance DisGeNET nanopublications discoverability in the Web of data. The general topic of DisGeNET nanopublications is ‘gene-disease association’, and each nanopublication is annotated using the Dublin Core vocabulary (DC) [20], object property ‘subject’, to state general topic, and the SIO concept ‘SIO_000983’ to state ‘gene-disease association’ as attribute.

**Fig. 1:**
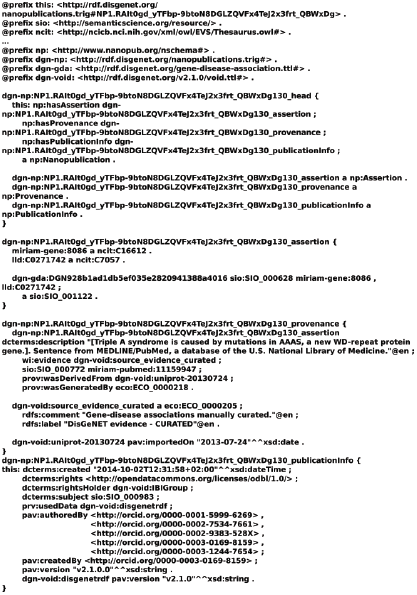
A DisGeNET nanopublication example in RDF/TriG notation.

This is a Linked Dataset as all resources are identified and linked to other Linked Datasets using de-referenceable URIs, and the most queryable entities such as gene and disease entities are identified by URIs constructed on widely adopted namespaces, such as the *identifiers.org* [21] to enable interoperability and query federation. In addition, the nanopublication URIs are *trusty*, i.e. they are generated by applying the Trusty URIs approach in order to guarantee the integrity of the nanopublication content and ensure its immutability, reliability, and verifiability for scientific citation.

### 3.1 Ontologies

The first step of modeling is to set specifically what and how the information has to be formally represented, and which ontologies to use to best represent the semantics. To represent the GDA in the assertion, we use the same triples and ontologies used in the RDF version. Therefore, we use SIO to encode both the type of association and to relate the disease and the gene associated, and the NCIt to encode the gene and disease biomedical entity types.

One important step in the nanopublication modeling was to establish which vocabularies were needed for the description of the provenance and metadata in the nanopublication graphs. To represent provenance information we mainly used the PROV Ontology (PROV-O) [22], which is a W3C recommended standard to core model the provenance, since it provides clear upper level classes, relationships and restrictions to frame any kind of provenance. For authorship, versioning, content creation information we used the Provenance, Authoring and Versioning (PAV) vocabulary [23], and for Web data we used the Provenance Vocabulary Core Ontology Specification (PRV) [24] as they are extensions of the PROV ontology that allow more explicit representations of the data. Finally, for general metadata the DC terms fit our requirements. The evidence annotation is formally well described using the Weighted Interests vocabulary (WI) [25], which provides a suitable object property *wi:evidence* that links the assertion with its evidence, and the Evidence Codes Ontology (ECO) [26] that provides suitable classes to term the meaning of the DisGeNET evidence classes (Table 1). It is not always straightforward to find ontologies that have classes that meet our requirements both, semantically and formally speaking (it was not possible to find an exact equivalence between our evidence classes and ECO classes). Sometimes, it may be really difficult but engines such as the NCBO Bioportal [27], Ontology Lookup Service [28] or tools such as NCBO Annotator [27] ease this task. Even though, the modeling in the Semantic Web offers the freedom to make any decision necessary to best describe the data, it is a good practice to use commonly used vocabularies. With this in mind, we chose the ECO and WI ontologies since they are already in use to model neXtProt evidence.

A summary of the Linked Data vocabularies used in DisGeNET nanopublications is shown in Table 2. We published our de-referenceable nanopublications on the Web with a human-readable list of the vocabularies used. Also, SIO and ECO are deployed in our triple store to be available both as machine-readable explicitly at axiom level to optimize the GDAs searches in our SPARQL endpoint, and to be human-readable in our Linked Data faceted browser.

**Tabel 2:**
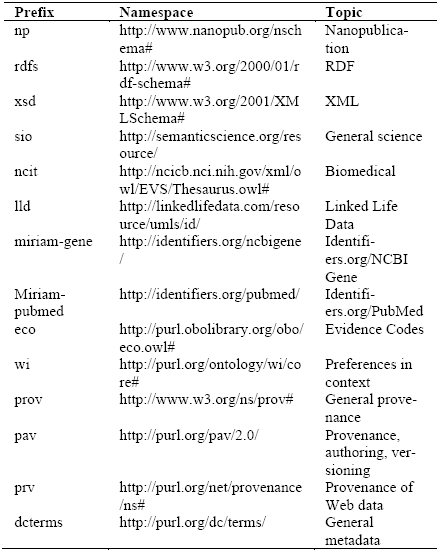
Vocabulary prefixes, namespaces, name and description of the nanopublications dataset.

### 3.2 Schema

Here we present the first implementation of the nanopublication model to DisGeNET that may change with future internal needs or external recommended adoption of standards. The official guidelines to create nanopublications accessed during summer of 2014 were used [29]. It is worth mentioning that the access to other nanopublications used as templates [13], neXtProt nanopublications [30], BEL2nanopub approach [31] facilitated the process of modeling. Moreover, the reuse of templates fosters a natural creation of standards both for RDF triple description of information and for vocabularies adoption and, consequently, enhances interoperability of scientific results [32].

The *Assertion graph* states the gene and the disease involved in the association, identified by URIs already used by the Semantic Web community to enhance interoperability. It also states the type of association asserted by the authors of the assertion using SIO which provides formal semantics for representing GDA entities, including the possibility to explicitly model the relationships between the gene and the disease in the association such as ‘genedisease association linked with altered gene expression’. The *Provenance graph* includes provenance and attribution information directly linked to the assertion such as the scientific article as the primary evidence, the source database from which was derived and the derivation date as the first stage of annotation. The method of extraction of the assertion is also included, i.e. if the statement was manually or automatically asserted. The level of evidence of each assertion is annotated using the DisGeNET evidence classification. This is an evidence classification system used in the context of DisGeNET, like it also exists in other databases such as the quality assessment classification system in neXtProt [30]. We minted the DisGeNET evidence concepts using as a base the DisGeNET namespace (http://rdf.disgenet.org). This is a temporary solution as, in order to make more discoverable DisGeNET nanopublications for aggregation of evidence, we are planning to use the ConceptWiki [33] to add URLs to the DisGeNET evidence concepts. Additionally, a human readable description of the assertion extracted from the attributed sources is added. Finally, the *Publication Information graph* includes all the metadata information regarding the nanopublication itself such as when it was made by date/time stamp, copyright information to inform how the nanopublication can be reused, the version of the nanopublication, the RDF Linked Data version used to produce it, and the authors and creators of the nanopublication. These attribution statements are done by object properties thanks to the unique digital identifiers linked to each researcher that can be obtained, e.g. ORCID identifiers.

### 3.3. Metrics, Versioning, Licensing

This is the first release of DisGeNET published as nanopublications and corresponds to the version v2.1.0.0. The dataset consists in 940,034 nanopublications, representing 940,034 scientific claims for 381,056 different GDAs with its detailed provenance, level of evidence and publication information description, all annotated as RDF statements and encapsulated into the nanopublication RDF graphs (3,760,136 graphs in total). Specifically, the dataset is composed of 31,961,156 N-Quads, i.e. RDF triples with its graph or context added as the fourth member in the tuple (Subject, Predicate, Object, Context), that are serialized in TriG syntax. The versioning track for nanopublications is conceived for recognition at both computer and human level. It consists of keeping track of the version’s provenance of both for the RDF and so for the relational version of DisGeNET, from which the RDF is derived. Thus, nanopublication versioning is a composition of versions: the version of the relational database (v2.1) + the version of the RDF dataset (v2.1.0) + version of the nanopublication (v0). Thus, v2.1.0.0 shows that is the first release of nanopublications with a zero number in the last position. The third position is reserved for the version of the RDF, a zero which states that is the first RDF derivation from the relational database. The version of the relational database is reserved for the first positions: v2.1. This dataset is made available as open under the Open Database License terms whose full text can be found at http://opendatacommons.org/licenses/odbl/1.0/.

## 4. Querying DisGeNET Nanopublications

With the aim to show the questions that can be answered by our nanopublication implementation, we use as example the following question: *What are the proteins (and their protein interactions) associated to Prostatic Neoplasms with curated evidence?* (Full query at DisGeNET Web site)

### 4.1. Retrieving Gene-Disease Associations

First, we query DisGeNET for all the genes associated to *Prostatic Neoplasms* (umls:C0033578). This query only involves the assertion graph.

~~~
SELECT DISTINCT ?gene WHERE {
?assertion a np:Assertion .
GRAPH ?assertion {
?gda sio:SIO_000628 ?gene,?disease .
?gene rdf:type ncit:C16612 .
?disease rdf:type ncit:C7057 .
filter regex(?disease, “umls/id/C0033578”)
}
} LIMIT 20
~~~

### 4.2. Filtering By Evidence

Second, we filter the prior results with those assertions annotated by ‘CURATED’ DisGeNET evidence. This query involves the provenance graph.

~~~
SELECT ?gene ?evidence WHERE {
?assertion a np:Assertion .
?provenance a np:Provenance . GRAPH ?assertion {
?gda sio:SIO_000628 ?gene,?disease .
?gene rdf:type ncit:C16612 .
?disease rdf:type ncit:C7057 .
filter regex(?disease, “umls/id/C0033578”)
}
GRAPH ?provenance {
?assertion wi:evidence ?evidence .
filter regex(?evidence, “curated”)
}
} LIMIT 20
~~~

### 4.3. Linking with Other LOD Resources

Finally, we cross DisGeNET prior results with the Interaction Reference Index database, which contains PPI annotations, through Bio2RDF::irefindex SPARQL endpoint, federating the query.

~~~
?gene sio:SIO_010078 ?protein_dgn . filter regex(?gene,"bio2rdf/uniprot”)
# Getting the interactome data from Bio2RDF::irefindex
SERVICE <http://irefindex.bio2rdf.org/sparql> {
optional{
?ppi a bio2rdf-ifx:Pairwise-Interaction ;
bio2rdf-ifx:interactor_a ?protein_dgn ;
bio2rdf-ifx:interactor_b ?protein_irx .
}
}LIMIT 20
~~~

Since in DisGeNET RDF is also represented the relation between gene and the protein/s that encodes, we are able to cross DisGeNET with Bio2RDF::irefindex by *Protein* resources through the corresponding linkset to <http://bio2rdf.org/uniprot:uniprotid>.

## 5. Availability, Production and Sustainability

DisGeNET nanopublications can be accessed in two ways: 1) Downloadable file in TriG format; 2) Queryable at the DisGeNET SPARQL endpoint (acces at http://rdf.disgenet.org/). The nanopublication production started from the relational database whose data is used to produce the RDF Linked Dataset. This RDF is generated by preprocessing in-house scripts that prepare data for the D2RQ platform [34], which is the software that serializes relational data into RDF/Turtle. Nanopublications Linked Dataset is derived from the RDF graph Linked Dataset representation. Both DisGeNET Linked Data are stored and available to query in the Virtuoso [35] SPARQL endpoint. The DisGeNET nanopublications should be able to be generated via SPARQL INSERT/CONSTRUCT queries over the RDF dataset. This is not currently possible for technical limitations of the Virtuoso SPARQL server. To bypass this temporary issue of the server, we developed custom scripts to construct and serialized in the recommended TriG syntax our nanopublications directly from our relational data. While, in one hand, this solution may be a source of production errors, it may foster a future base of scripts for a sustainable pipeline production of DisGeNET nanopublications on a regular basis and shareable to the community through systems as GitHub.

## 6. Applications

One of the future goals of DisGeNET is to be integrated with the Open PHACTS Discovery platform. The Open Pharmacological Concepts Triple Store project (Open PHACTS) has developed a powerful an open access, cloud-based, innovation data platform via a Semantic Web approach that allows scientists to draw on diverse databases to answer all kinds of questions relating to drug development. The new version of the platform will integrate and provide access to additional datasets such as WikiPathways [4], neXtProt, which were also recently converted into nanopublications, or DisGeNET. This will be a real test for the integration of DisGeNET along with other sources, primarily as RDF and then as nanopublications, that can provide experience about it to the Semantic Web community.

## 7. Summary

We have created a nanopublication-based Linked Dataset that provides 940,034 nanopublications about scientific statements of human GDAs. These GDAs published as Trusty URI nanopublications are machine-interpretable, immutable, perenne, and verifiable to promote citation. Each GDA statement has its provenance description providing attribution, how, when and from where it was generated to confer trust. Each GDA is classified as “CURATED”, “PREDICTED”, or “LITERATURE” in the Dis-GeNET context to categorize the evidence of the statement based on the type of assertion and curation made in the original databases. We recommend enriching the provenance annotation stating the type of curation of the assertion, and if it is possible classify the nanopublications by its level of evidence. Dis-GeNET nanopublications include metadata annotation about the general topic of the nanopublication, i.e. ‘Gene-Disease Association’, semantically described by SIO to ease its discoverability in the Semantic Web. With illustrative queries we show how to explore GDAs with DisGeNET nanopublications and how to integrate them with relationships published in other LOD sources. The publication on the Web of data of DisGeNET nanopublications will enable a large-scale interconnection of statements, even crossing domains, to explore them based on evidence. This is essential for discovery science, and with the nanopublishing of DisGeNET we aim to provide a better picture of the molecular basis of pathological conditions.

## Acknowledgments

We thank Dr. Mark Thompson (Leiden University Medical Center) and Dr. Jesse Van Dam (Wageningen UR) for their expertise in the modeling of nanopublications. Funding: The research leading to these results has received support from Instituto de Salud Carlos III-Fondo Europeo de Desarollo Regional (PI13/00082 and CP10/00524), the Innovative Medicines Initiative Joint Undertaking under grants agreements n° 115002 (eTOX) and n° 115191 (Open PHACTS)], resources of which are composed of financial contribution from the European Union’s Seventh Framework Programme (FP7/2007-2013) and EFPIA companies’ in kind contribution. Laura I. Furlong received support from Instituto de Salud Carlos III Fondo Europeo de Desarollo Regional (CP10/00524). The Research Programme on Biomedical Informatics (GRIB) is a node of the Spanish National Institute of Bioinformatics (INB).

